## Supplementary Files for "TuNR: Orthogonal Control of Mean and Variability of Endogenous Genes in a Human Cell Line"

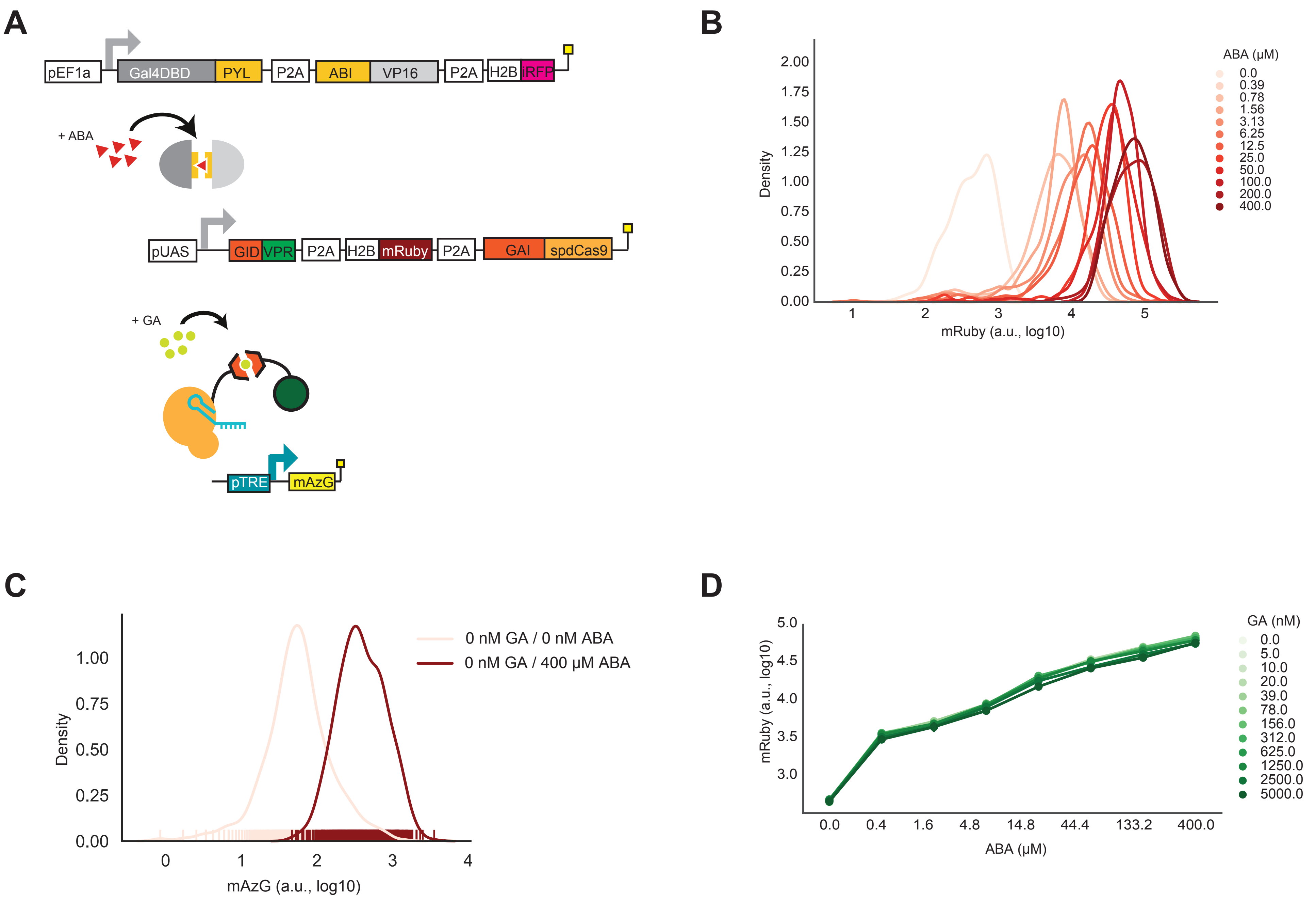

**Supplementary Figure 1**

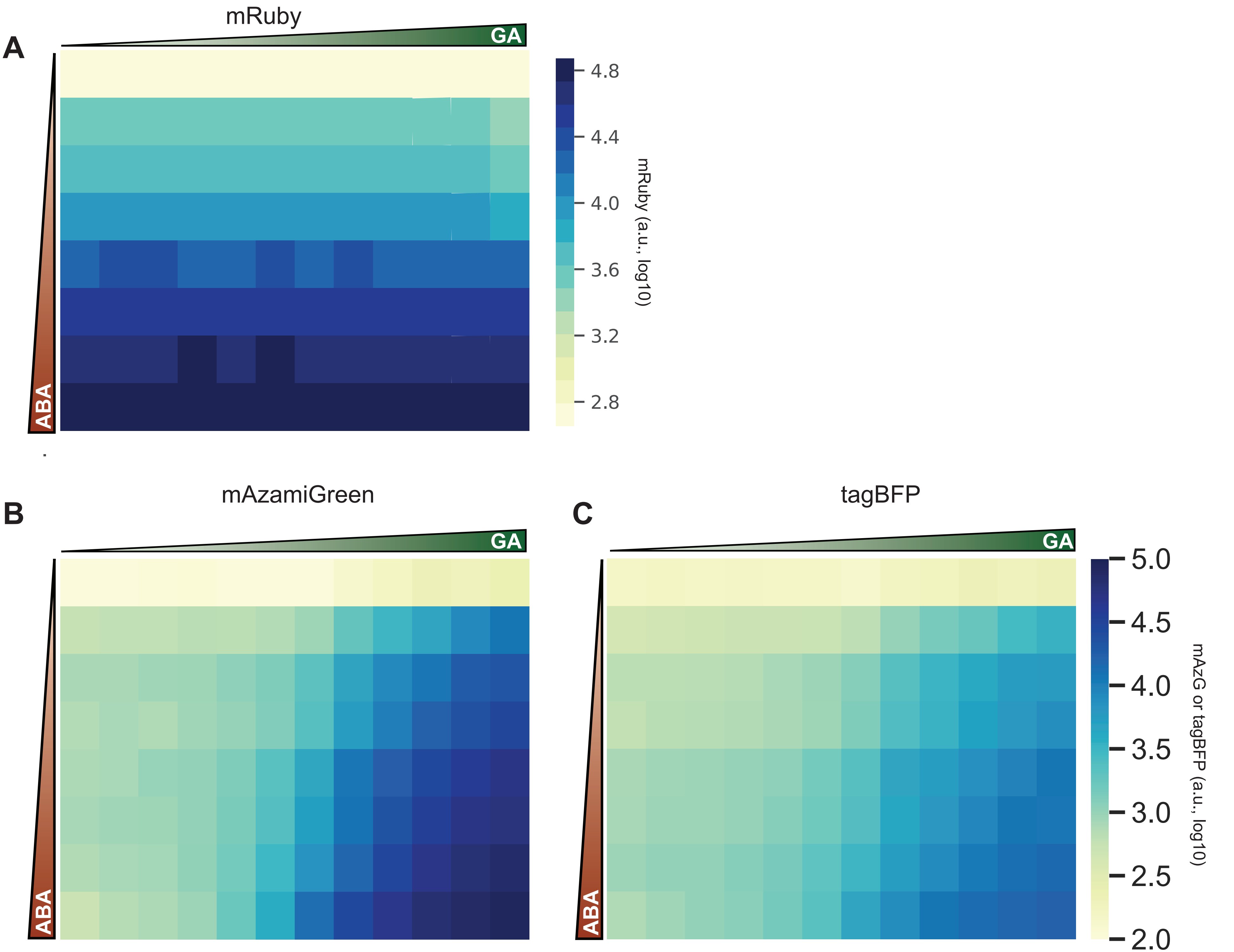

**Supplementary Figure 2**

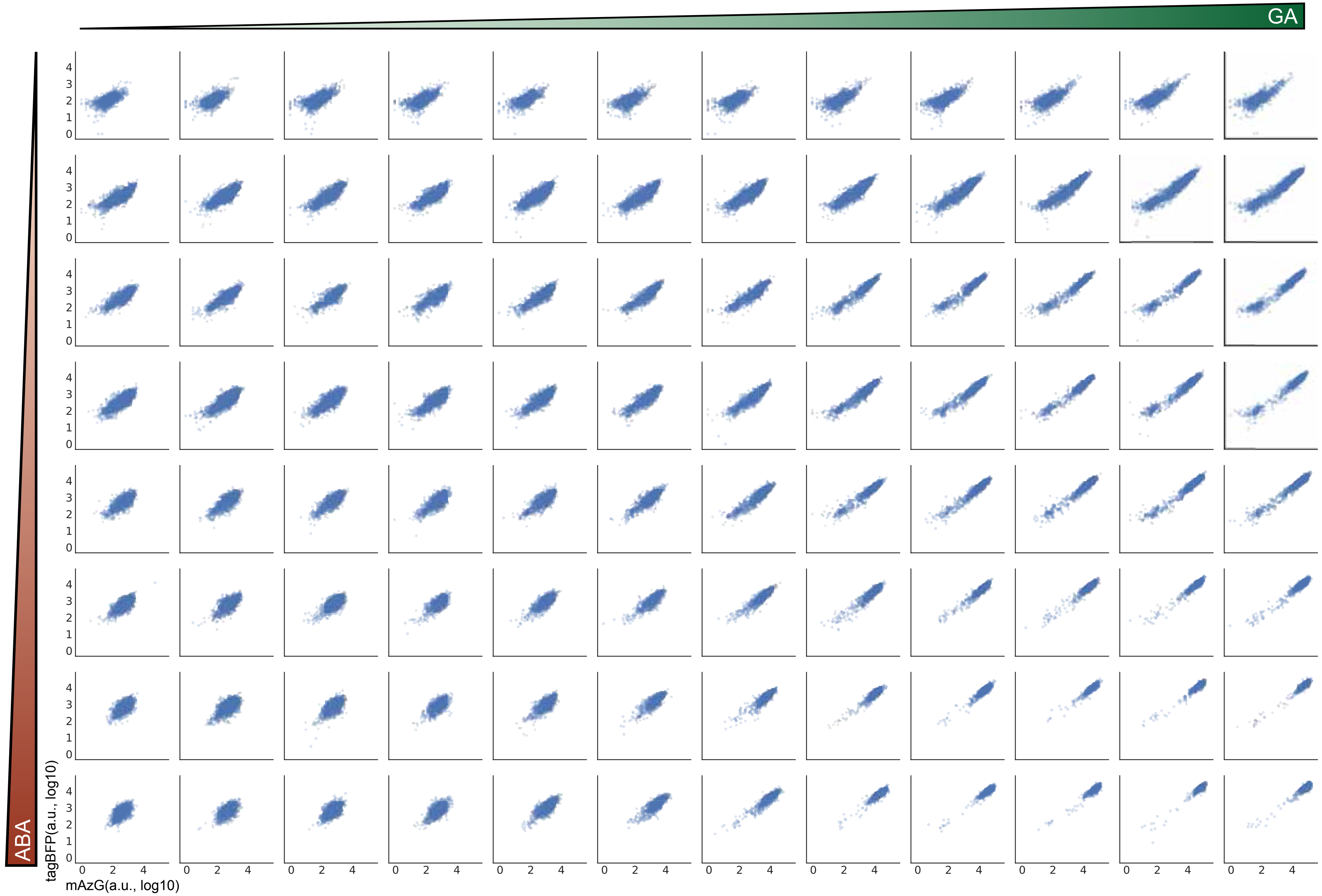

### Supplementary Figure 3

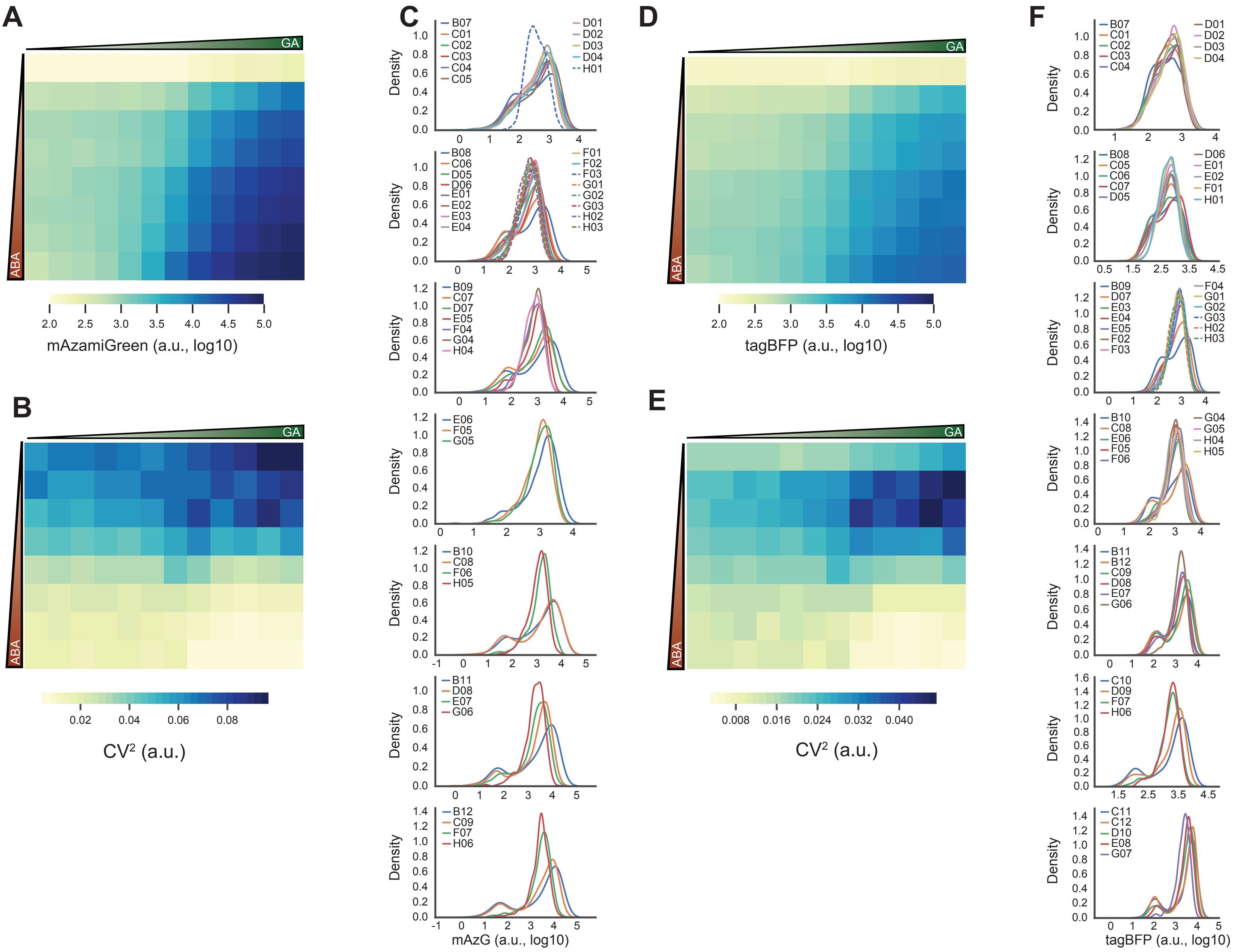

**Supplementary Figure 4**

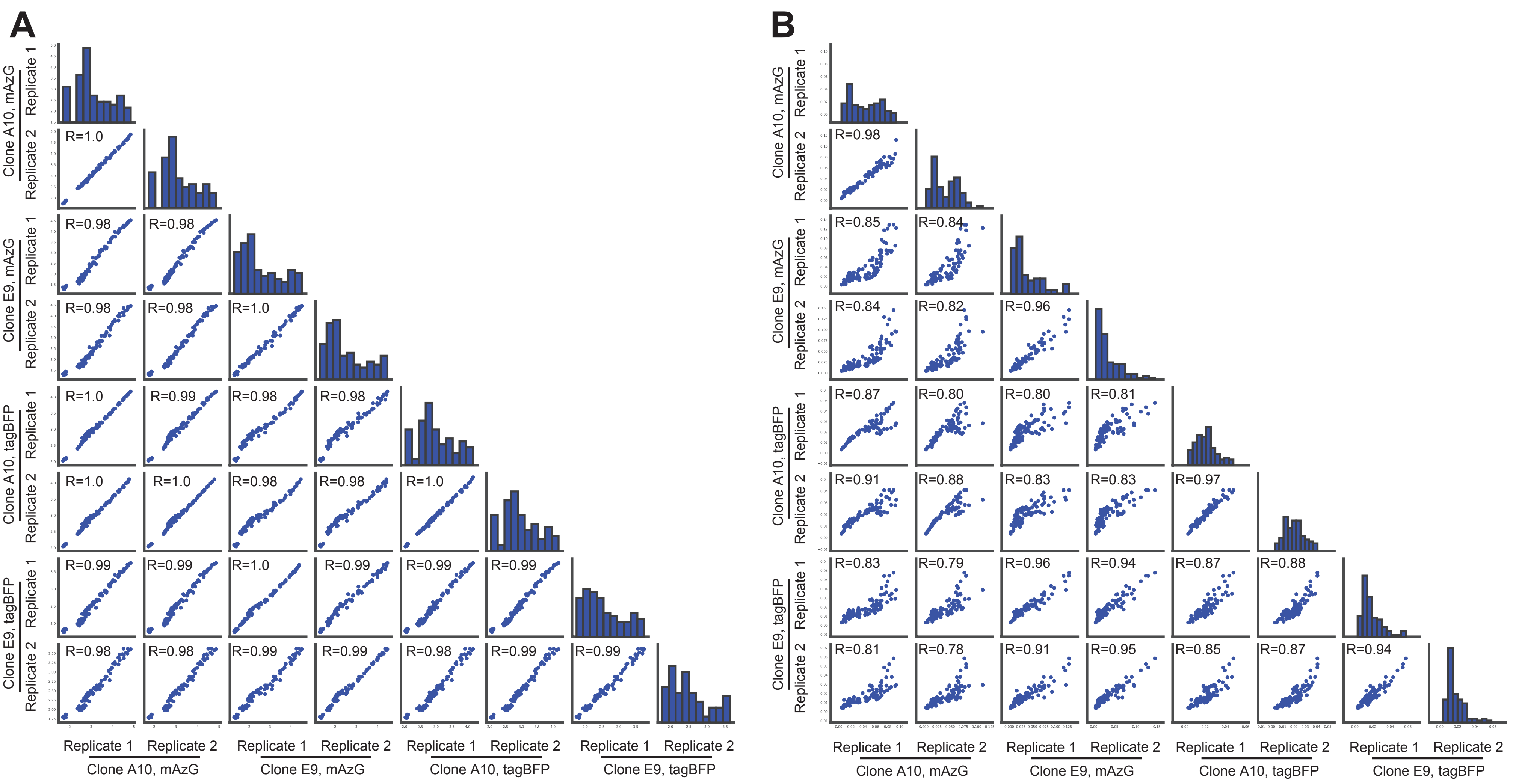

**Supplementary Figure 5**

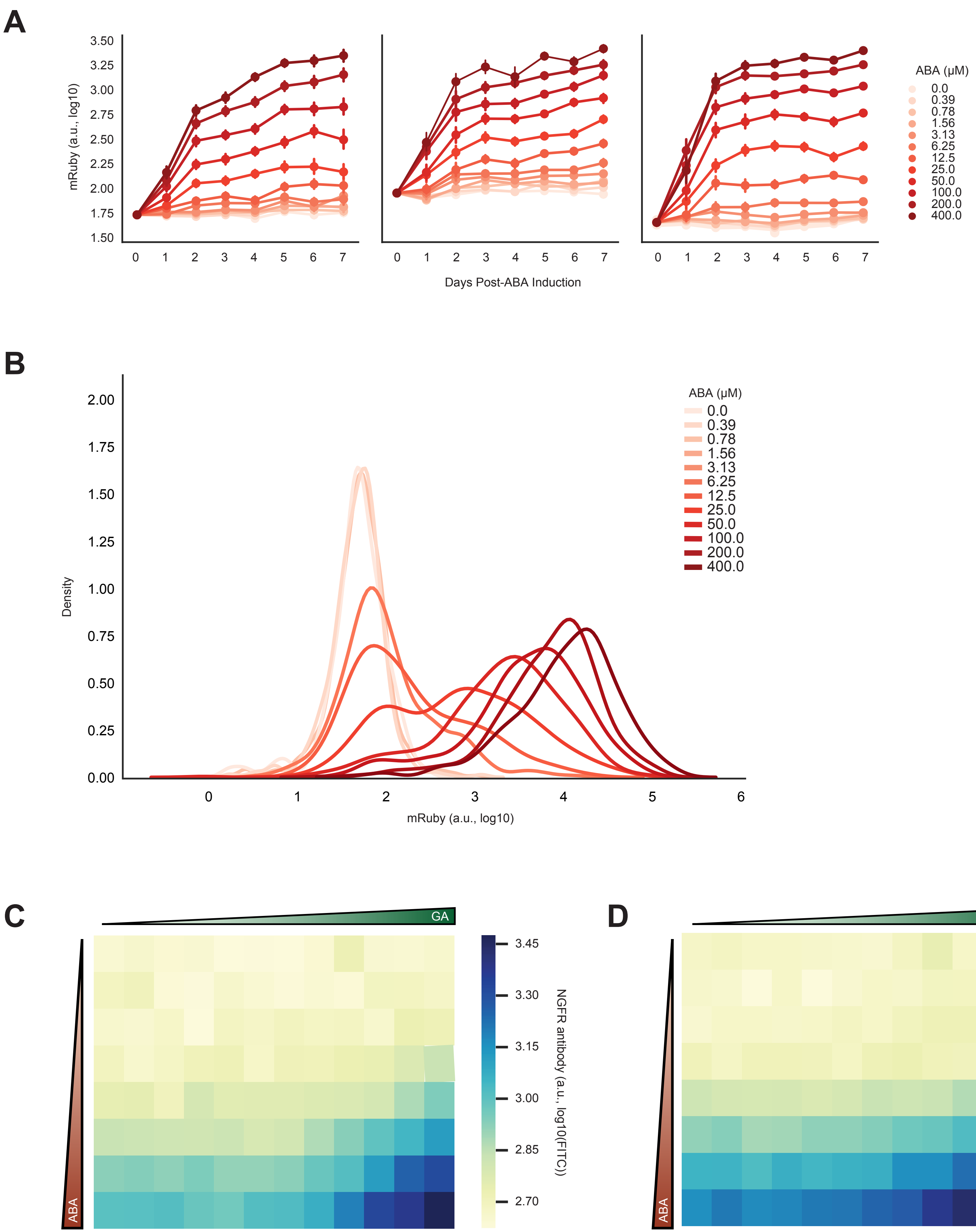

**Supplementary Figure 6**

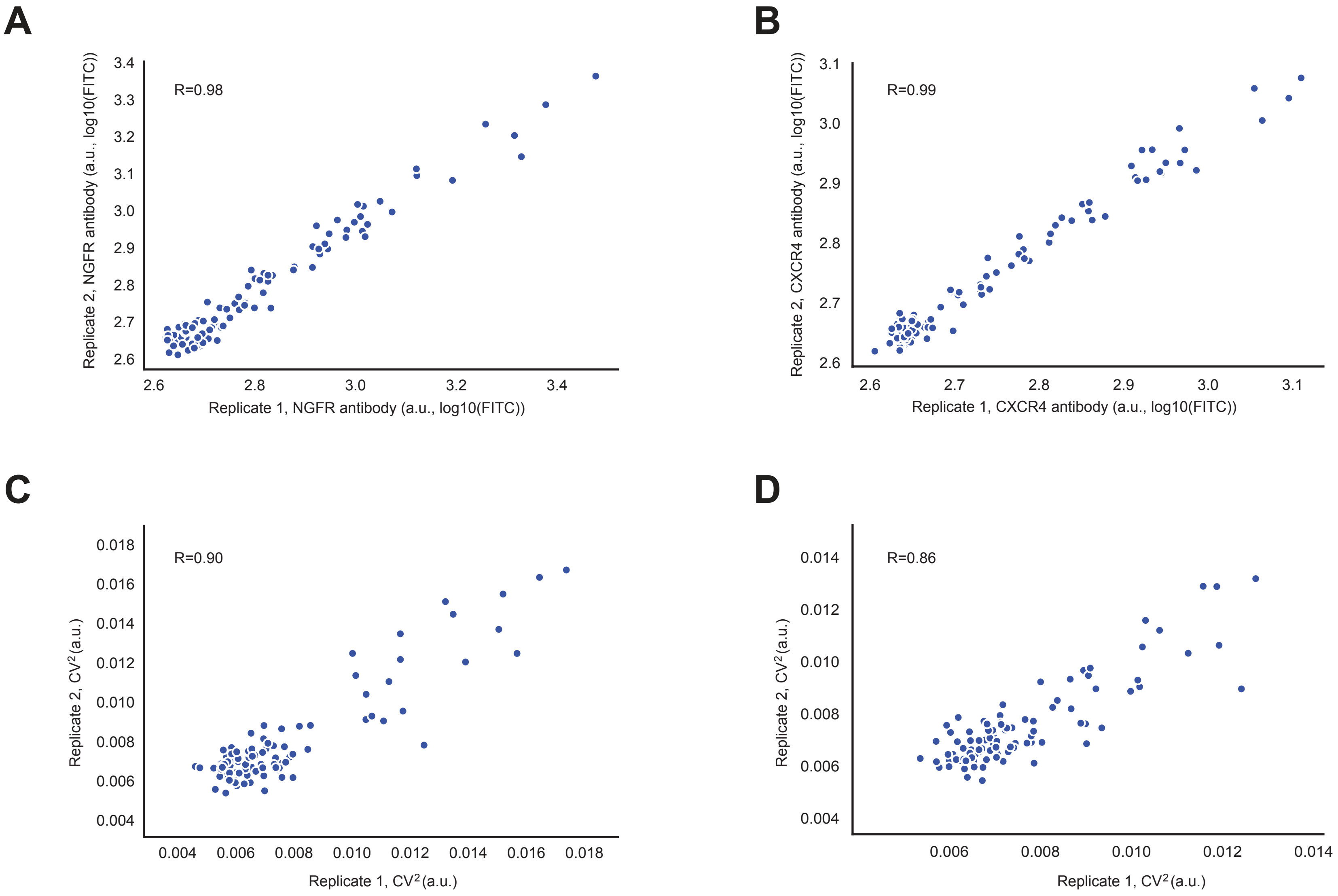

**Supplementary Figure 7**

| Figure | Description | Plasmid Name | Notes |
|---|---|---|---|
| 1,2 | Constitutive split Gal4-VP16, iRFP | ARB298 | TuNR Part 1 |
| 1,2 | pUAS-split dCas9-VPR, mRuby | ARB255 | TuNR Part 2 |
| 1,2 | pTRE-GFP,BFP, mu6-gRNA | ARB256 | TuNR Part 3 |
| 3 | Constitutive split Gal4-VP16 | ARB298 | TuNR Part 1 |
| 3 | pUAS-split dCas9-VPR | ARB255 | TuNR Part 2 |
| 3 | mu6-CXCR4gRNA, pEF1a-BFP | ARB498 | CXCR4 gRNA Lentivirus |
| 3 | mu6-NGFRgRNA, pEF1a-BFP | ARB499 | NGFR 3xgRNA Lentivirus |
| n/a | Full TuNR Circuit | ARB293 | Full TuNR on one plasmid |
| n/a | Chassis TuNR Circuit | ARB461 | TuNR on one plasmid minus gRNA, GFP, BFP |

Supplementary Table 1

| Figure | Target Locus | Protospacer Sequence | Backbone Vector | Notes: |
|---|---|---|---|---|
| 1,2 | pTRE | TACGTTCTCTATCACTGATA | MTK234_002 | mu6 promoter, stem region 1 |
| 3 | CXCR4_1 | GGGGCAGACGCGAGGAAGGA | MTK234_002 | mu6 promoter, stem region 1 |
| 3 | NGFR_1 | GccgcccTGCCGCCTCGGGG | MTK234_002 | mu6 promoter, stem region 1 |
| 3 | NGFR_2 | GTGAGAGGCTCTAAGGGACA | MTK234_055 | bu6 promoter, stem region 2 |
| 3 | NGFR_3 | GCGGGCGGGCGCGGTTCCGG | MTK234_056 | hu6 promoter, stem region 3 |
| General Notes: | | | | |
| gRNAs were oligo-annealed into backbone vectors and further subcloned into lentiviral vectors | | | | |
| 20N corresponds to Protospacer Sequence | | | | |
| MTK234_002 | 5' - TGTTTG - 20N - G - 3' | | | |
| | 5' - TAAAC - 20 N(Reverse Complement) - CA - 3' | | | |
| MTK234_055 | 5' TTATG-20N-G 3' | | | |
| | 5' CAAAC-20N (Reverse Complement) - C 3' | | | |
| MTK234_056 | 5' ACATG-20N-G 3' | | | |
| | 5' GAAAC-20N (Reverse Complement) -C 3' | | | |

Supplementary Table 2
